# Antitumor effects of Tv1 venom peptide in liver cancer

**DOI:** 10.1101/518340

**Authors:** Prachi Anand, Petr Filipenko, Jeannette Huaman, Michael Lyudmer, Marouf Hossain, Carolina Santamaria, Kelly Huang, Olorunseun O. Ogunwobi, Mandë Holford

## Abstract

A strategy for treating the most common type of liver cancer, hepatocellular carcinoma (HCC) applies a targeted therapy using venom peptides that are selective for ion channels and transporters overexpressed in tumor cells. Here, we report selective anti-HCC properties of Tv1, a venom peptide from the predatory marine terebrid snail, *Terebra variegata.* Tv1 was applied *in vitro* to liver cancer cells and administered *in vivo* to allograft tumor mouse models. Tv1 inhibited the proliferation of murine HCC cells via calcium dependent apoptosis resulting from down-regulation of the cyclooxygenase-2 (COX-2) pathway. Additionally, tumor sizes were significantly reduced in Tv1-treated syngeneic tumor-bearing mice. Tv1’s mechanism of action involves binding to specific transient receptor potential (TRP) cation channels that are overexpressed in HCC cell models. Our findings demonstrate the unique potential of venom peptides to function as tumor specific ligands in the quest for targeted cancer therapies.

---

The second most common cause of cancer-related death worldwide is hepatocellular carcinoma (HCC), which is the most common form of liver cancer.^1^ Despite liver cancer’s growing prevalence, there is a startling lack of treatments and currently only one small molecule, Sorafenib (Nexavar^®^), is approved by the U.S, Food and Drug Administration to treat advanced HCC. Venom peptides from predatory animals exhibit extraordinary specificity and selectivity for cancer cells and could be the turnkey for addressing the unmet need of identifying new agents for treating HCC.^2–5^ Whether by acting directly as antitumor therapeutics, diagnostic tags, adjuvants or just carriers for other relevant moieties, venom peptides or their derived conjugates have shown significant potential as bioactive agents that can overcome the limitations of developing tumor specific cancer therapies.^2,4,6^

Specifically, venom peptides have been shown to mediate anticancer activity via interactions at tumor cell membranes and as intracellular components affecting cell migration, invasion, and angiogenesis—all major functional mechanisms of tumor growth. The RGD (Arg-Gly-Asp) consensus sequence from the disintegrin class of venom peptides and NGR (Asn-Gly-Arg) have been used as scaffolds to develop anti-metastatic drugs, such as Cilengitide^®^.^7–9^ Some venom peptides, such as chlorotoxin (CTX) isolated from *Leiurus quinquestriatus* scorpion venom, display efficient tissue penetration and uptake by heterogeneous cranial cancer tissues. CTX has led to the development of several theranostic brain tumor imaging drugs (BLZ-100 and TM601) that are referred to as tumor paint and used to localize glioma cells.^10,11^ Venom peptides have also been chimerized with existing chemotherapeutics, and functionalized as carrier vehicles for drugs with lower selectivity or bioavailability.^12^

Two recent examples of conoidean marine snail venom peptides that identify or inhibit specific ion channels and are also related to cancer are ziconotide and k-PVIIA. Ziconotide (Prialt^®^), discovered from the venomous marine snail *Conus magus*, is a 25 amino acid peptide (MVIIA) used to alleviate chronic pain in HIV and cancer patients by inhibiting N-type calcium channels.^13^ Alternatively, k-PVIIA peptide from *Conus purpurascens*, selectively blocks the voltage-gated Shaker potassium (K^+^) channel, and was found to mediate tumor cell proliferation by binding to hERG, a K+ channel protein that increases in concentration on the cell surface of cancer cells.^14^ Taken together, the antitumor activity of venom peptides RGD, CTX, MVIIA, and k-PVIIA is a persuasive argument for how ion channels and transporters can be effective new molecular targets for cancer therapies.

This is confirmed by recent compelling experimental evidence that pharmacological inhibition of ion channels or their regulators counteracts tumor growth, prevents metastasis and overcomes therapy resistance of tumor cells.^15–17^

Metastasis, the main cause of cancer-associated mortality, depends on two key processes: i. cell migration of cancer cell to invade adjacent tissues followed by intravasation into blood/lymphatic vessels and ii. tumor vascularization, which gives access to the blood stream. Cell migration and tumor vascularization are often associated with changes in ion channel expression and/or activity. In particular, Ca^2+^ channels are of importance because Ca^2+^ is the key messenger regulating signaling pathways in cellular processes such as proliferation, apoptosis, transcription, migration and angiogenesis.^18,19^ In this context, the recently identified Ca^2+^ channel family, Transient Receptor Potential (TRP), has been associated with several cancers and its role has been increasingly clarified over the last two decades.^20,21^

TRP channels modulate intracellular Ca^2+^ concentrations, controlling critical cytosolic and nuclear events that are involved in cancer initiation and progression. Therefore, it is anticipated, that the expression and function of some TRP channels is altered during tumor growth and metastasis.^22^ Recent reports suggest the expression and/or activity of TRP channels mark and regulate specific stages of cancer progression.^21,23,24^ As such, TRP channels can be envisioned as polymodal molecular sensors suggesting that the physiological relevant stimulus for any given TRP will be governed by the specific cellular context (i.e. phosphorylation or dephosphorylation, lipid environment, interacting adjacent proteins and concentration of related ligands), which dramatically changes during carcinogenesis. Among the TRP channel families, TRPCs, TRPMs and TRPVs are mainly related to malignant growth and progression. In particular, TRPC6 and TRPV6 have recently been reported to play a critical role in the development of many carcinomas including human renal cell carcinoma^25^, prostate cancer^21^, lung cancer,^26^ and other types of cancer.^23,27–30^ Studies of TRP protein expression in liver tumor cell lines also suggest that altered expression/function of TRPC6 and other TRP channels may play a role in the development, progression, and metastasis of HCC.^31^

Here, we present the anticancer and anti-tumorigenic properties of recently identified venom peptide Tv1, from predatory marine snail *Terebra variegata* (Fig. 1). Tv1, a 21 amino acid peptide with unique structural properties compared with known snail venom peptides.^32^ Tv1 was chemically synthesized and assayed using both *in vitro* and *in vivo* systems. Our results suggest that Tv1 inhibits HCC selectively, and that its mechanism of action involves downstream manipulation of TRPC6 and/or TRPV6 channel activity, which were overexpressed in the HCC models used in this study.

**Figure 1.**
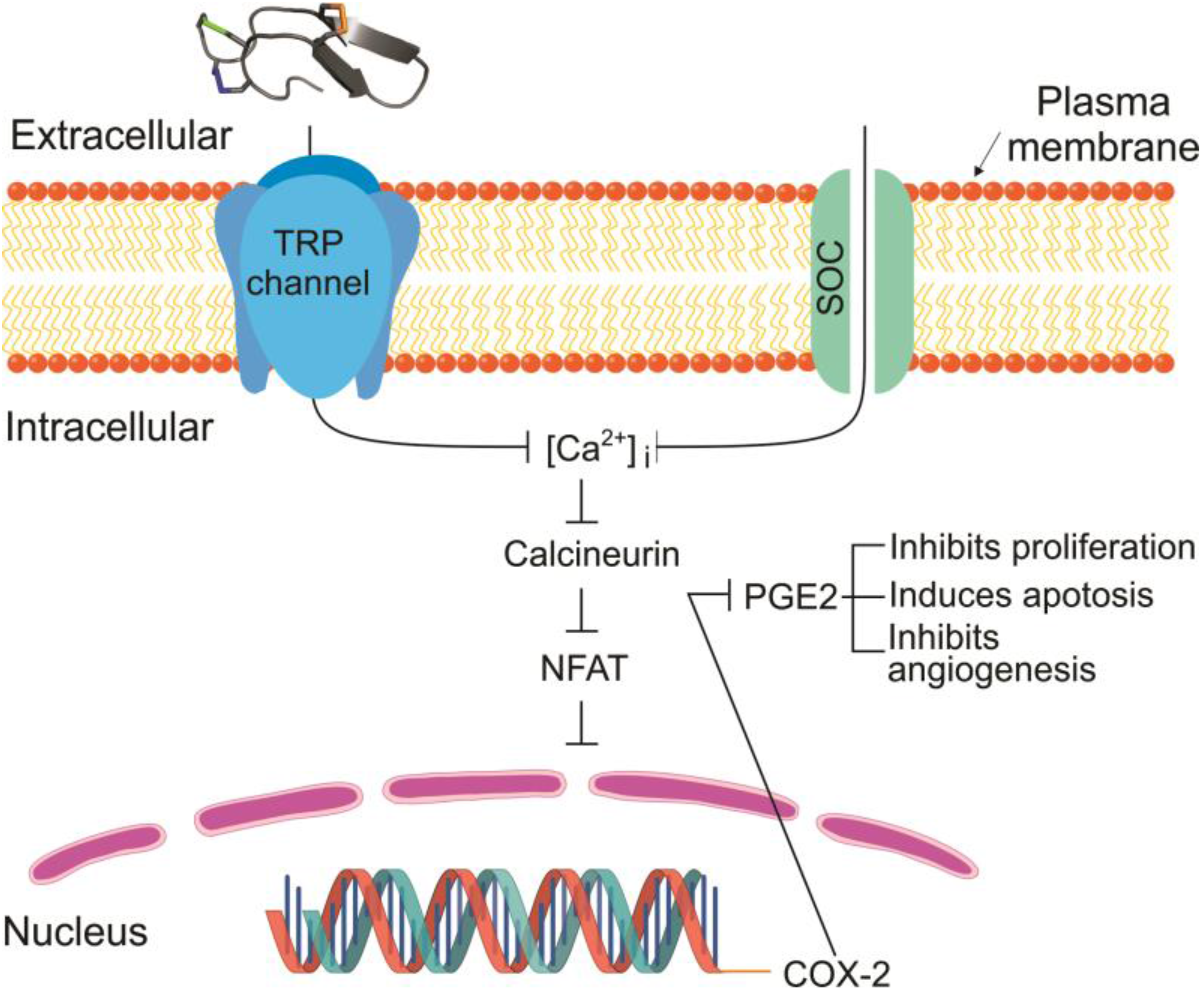
Potential mechanism of Tv1 antitumor activity inhibits COX2 and PGE2 function. In our model of Tv1 antitumor activity in liver cancer cells, overexpression of TRP channels (TRPC and V6) stimulate COX-2-dependent PGE2 production via enhanced [Ca^2+^] dynamics. Influx of Ca^2+^ can occur through voltage gated (VGC), receptor operated (ROC), and store operated (SOC) calcium channels. Transient receptor potential (TRP) channels contribute to store operated calcium (SOC) channels. Ca^2+^-dependent transcription factor NFAT is activated via dephosphorylation by calcineurin, which is activated upon binding of Ca^2+^/calmodulin. Ubiquitously present transcription factor NFAT regulates COX-2 expression and further prostaglandin E2 release in different cancer cells and its activation occurs through Ca^2+^ influx associated with TRPC1-, TRPC3-, or TRPC6-associated SOC or ROC activities. PGE2 release plays multiple roles in cancer as shown. Upon Tv1 inhibition of TRP channels all the downstream pathways leading to proliferation inhibition and apoptosis of tumor cells will encounter.

## Results

### Tv1 shows selective cytotoxicity in liver cancer cells

Cytotoxicity of Tv1 was tested using an MTT assay in cervical (HeLa), neuroblastoma (SKnSH), prostate (WPE1-NA22), and liver (BNL 1ME A.7R.1 (1MEA)) cancer cell lines. Cell viability was measured after 48 h of treatment with Tv1 and doxorubicin (Dox), a commercial anticancer drug that was used as a positive control. MTT analyses revealed that Tv1 displayed significant (p<0.01) cytotoxicity in 1MEA (liver cancer) cells compared with the other cancer cell lines tested (Fig. 2a). In contrast, Dox was cytotoxic (p<0.01) to all the cell lines used (Fig. 2a). An MTT cytotoxicity assay was also used to examine the comparative cytotoxicity of non-tumorigenic (BNL.CL.2 (BNL)) and tumor (1MEA) liver cells with Tv1, Dox and Sorafenib (Sora), a nonselective commercial liver cancer drug (Fig. 2b). Dox and Sora did not show any selectivity in toxicity for BNL (non-tumorigenic) and 1MEA (liver cancer) cells. In contrast, Tv1 was selective for tumor versus non-tumorigenic liver cells (Fig. 2b). Tv1 was also tested in the human liver cancer cell line HepG2 and the results indicate that at higher micromolar concentrations, Tv1 also has cytotoxic effects on HepG2 cells (Fig. S1).

**Figure 2.**
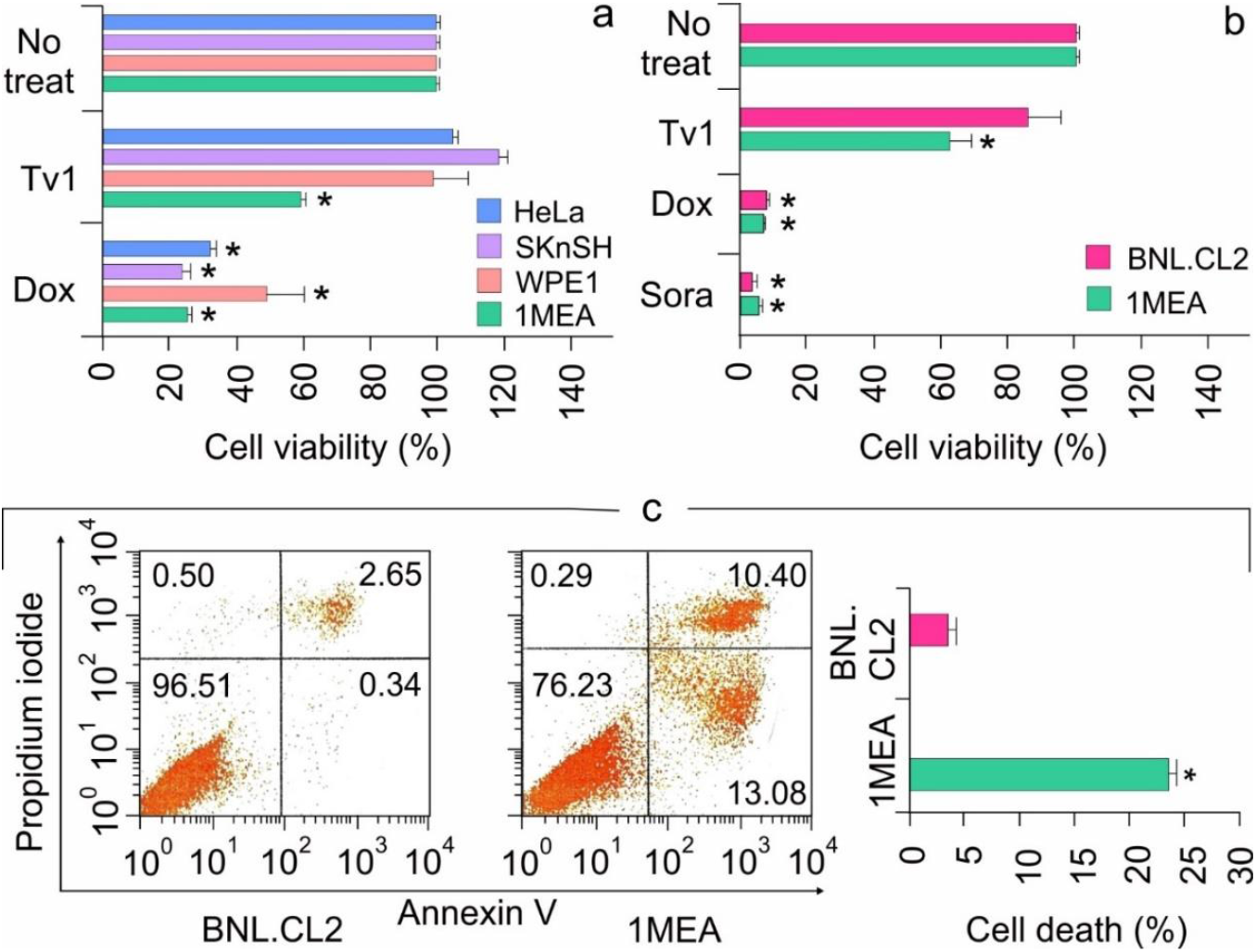
Tv1 selectively cytotoxic against liver cancer cells. **(a)** Cytotoxicity of Tv1 in different cancer cells. 48 h treatment of Tv1 showed its selective cytotoxicity for mouse liver cancer (1MEA) cells with a significant cell death of 40%. (n=5). Origin of cancer cells lines used: HeLa-cervical cancer, SKnSH-neuroblastoma cells, WPE1-prostate cancer, 1MEA-liver cancer. **(b)** Comparative cytotoxicity of Tv1 with anticancer drugs Doxorubucin and Sorafenib, which are commercially available. Tv1 demonstrated selective cytotoxicity to liver cancer cells, with diminished activity on normal liver cells, while Sorafenib and Doxorubicin showed high cytotoxicity and non-selectively in both the cell lines (n=3). (c) Flow cytometry using AnnexinV/PI staining resulted in significant apoptotic cell death by 16 h Tv1 treatment. n=3 *p<0.01.

### Tv1 induced apoptotic death of liver cancer cells and inhibited migration

The functional mechanisms modulated by Tv1 treatment of non-tumorigenic (BNL) and tumorigenic (IMEA) cells were examined using Annexin V/PI staining, a marker for apoptosis (Fig. 2c). BNL and 1MEA cells were treated with Tv1 for 16 h, and then were processed for staining. A total of 23% cell death was observed in 1MEA, compared to 3.1% in BNL cells (p<0.01). Of the 23% Tv1-induced cell death, >13% of the cells were stained with Annexin V, indicating that Tv1 treatment induced apoptotic cell death. Additionally, as cancer cell invasion and metastasis are characterized by epithelial-mesenchymal transition (EMT), we examined cell migration (BNL and 1MEA) using wound-healing assays. Within 6 h, compared with untreated 1MEA cells, there was a significant difference (p<0.001) in the wound size (cell covered area) of Tv1-treated 1MEA cells (Fig. S2). Treatment with Dox did not have a significant effect on wound healing.

### Tv1 has antitumor activity in an allograft tumor mouse model

To determine if the *in vitro* effects of Tv1 treatment would translate to an *in vivo* scenario, we used an allograft tumor mouse model in which 1MEA cells were implanted in Balb/c mice. Tv1 was intraperitoneally injected at a low dose (0.08 mg per kg body weight (mpk)) and at a high dose (0.8 mpk), daily for 8 d (days), and then after a break of 4 d, Tv1 was administered again for 7 d (Fig. 3a). In both low and high dose administered animals, Tv1 inhibited tumor growth without any evidence of systemic toxicity, as evidenced by normal body condition scores of both experimental and control animals. Tv1 at high dose caused a significant 47% reduction in tumor volume at the end of the observation period. In comparison, administration of control vehicle resulted in an average increase in tumor volume by 60% (from 323.74 mm^3^ to 796.8 mm^3^). Tv1’s reduction of tumor growth was statistically significant (p<0.01) as early as the third day after beginning treatment, suggesting rapid action of the peptide. Throughout the treatment period, gross tumor sizes were also compared among the no-treatment and treatment groups, and a highly significant difference (p<0.001) was found in tumor volumes of the no-treatment control versus high dose Tv1 treatment groups (Fig. 3b).

**Figure 3.**
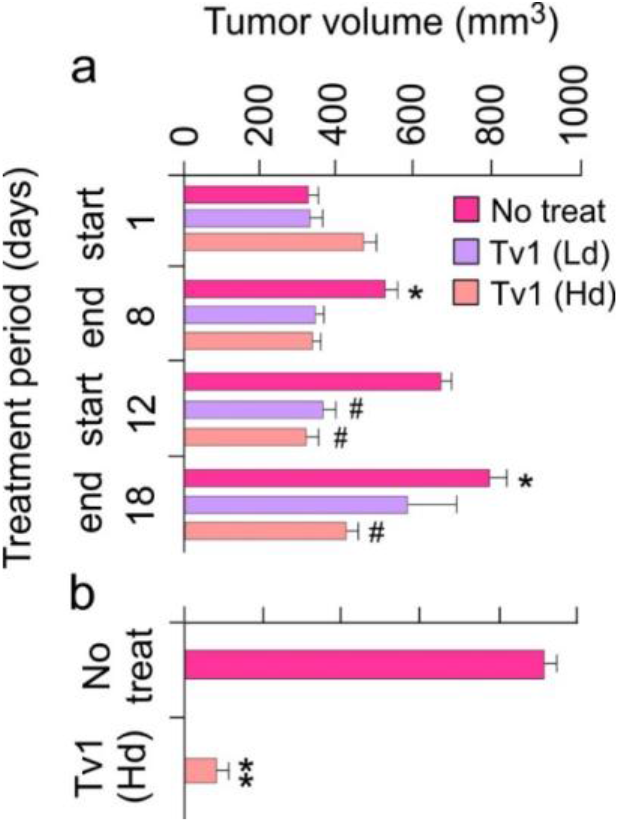
*In vivo* antitumor activity of Tv1 in tumor allograft mouse model. (a) Tumor volumes over a period of 18-day treatment showing significant reduction in tumor sizes in the first phase of treatment (1-8 day) in high (Hd-0.8 mg/kg body weight) and low (Ld-0.08 mg/Kg body weight) dose treated group and a highly significant increase in the tumor volumes in no treatment groups. * p<0.01 compared with day 1, #p<0.01 when compared with no treatment group at respective time. (b) In comparison of no treatment and Tv1 high dose treated tumor volumes throughout the treatment period, Tv1 treated mice had a significantly smaller tumor volume. **p<0.001 compared with no treatment group.

### TRPC6 and TRPV6 ion channels overexpressed in 1MEA cells

Based on their previously well-known role in cancer and metastasis, the presence of four ion channels (HERG/Kv11.1, TRPC1, TRPC6, & TRPV6) were screened in (BNL and 1MEA) cells using a high throughput 96-well in-cell western assay (ICW)^21,33,34^. Anti-TRPV6 and anti-TRPC6 channel antibodies specifically, and the cells in general, were observed by IR Fluorescence at 800 and 700 nm, respectively. TRPC6 and TRPV6 are overexpressed in 1MEA cells compared with BNL cells (Fig. 4a). Incubation with anti-HERG and anti-TRPC1 channel antibodies did not result in fluorescent signals, indicating their absence in both BNL and 1MEA cells (Fig. S3). To control for background and non-specificity, wells treated with specific antigens for antibodies showed no green fluorescence, meaning that the antibodies were specific to the respective ion channels, while the presence of red signal indicated the presence of cells. Of the channels tested, TRPC6 and TRPV6 channels were the most dominant channels present and overexpressed in 1MEA cells (Fig. 4a).

**Figure 4.**
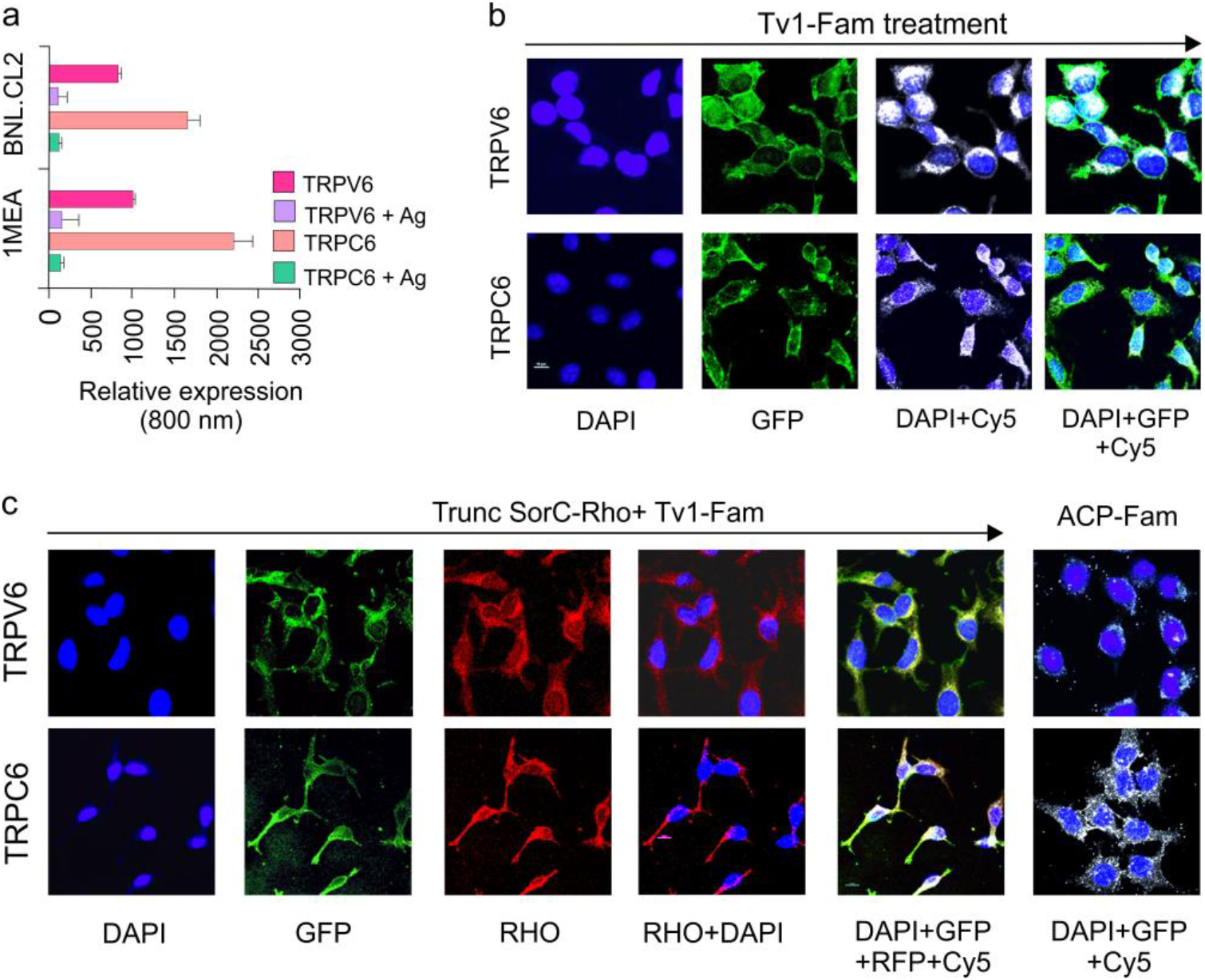
Tv1 co-localizes with TRP Channels. **(a)** Intensities are compared at 800nm channels after normalizing at 700nm channel for call-tag. The analysis demonstrated significant increased levels (p<0.05) of TRPV6 and TRPC6 in 1MEA cells compared to BNL cells. **(b)** Co-localization analysis of Tv1 with the ion channels TRPC6 and TRPV6 in 1MEA cells using MOC (manders overlap coefficient). y-axis showing the incubation with different probes, x-axis showing the filters used for imaging. High MOC values confirmed the Tv1 co-localizing with these channels. **(c)** A specific blocker for TRPV6 channel, truncSorC-Rho was used to study the colocalization of Tv1-Fam with TRPV6 channels in its presence. High MOC for truncSorC-Rho and Tv1-Fam with channels represent their colocalization. A fluorescently labeled non-specific peptide ACP-Fam, probed with channel antibodies to examine non-specific green fluorescence and no colocalization was detected. DAPI-staining nucleus for the presence of cells, GFP-green channel for the presence Tv1-Fam, Rho channel for the presence of Rhodamine labeled truncSorC-Rho peptide, Cy5-cy-5 channel for the presence of ion channels as secondary antibodies used were cy-5 labeled.

### Tv1 co-localizes with TRP channel subtypes

To examine if the observed anticancer activity of Tv1 was due to its activity on the TRPC6 and TRPV6 channels overexpressed in 1MEA cells, an oxidized and fluorescently-labeled Tv1 peptide (Tv1-Fam) was synthesized and used in an immunofluorescence assay (IFA) to determine colocalization of Tv1-Fam with TRPC6 and TRPV6. Cells were incubated with Tv1-Fam for 30 min prior to separate incubations for TRPC6 and TRPV6 IFAs]. Mander’s overlap coefficient (MOC) values were calculated for 3 independent experiments and a mean value of 0. 80±0.01 and 0.9±0.05 for TRPC6 and TRPV6, respectively, suggested that Tv1-Fam colocalized with TRPC6 and TRPV6 channels (Fig. 4b). In a similar assay as Tv1-Fam, we also tested the only reported peptide inhibitor for TRPV6, soricidin peptide (SorC) from the paralytic venom of northern short-tailed shrew *(Blarina brevicauda)^35^.* A truncated version of soricidin (truncSorC) was synthesized and labeled with Rhodamine (truncSorC-Rho) to perform IFA in the presence of Tv1-Fam for TRPV6 channels. The orthogonal labeling of Tv1-Fam and SorC-Rho enabled us to visualize both peptides colocalizing with TRPV6 channels.

TruncSorC-Rho co-localized with TRPV6 channels with a mean MOC of 0.88±0.04 (Fig. 4c). The truncSorC-Rho MOC value is similar to that of Tv1-Fam and TRPV6 (0.9±0.05). When Tv1-Fam and truncSorC-Rho were applied at the same time, both produced a MOC of 0.92±0.01, confirming that both Tv1-Fam and truncSorC-Rho were co-localized with TRPV6 ion channels. To further confirm if the specific binding of Tv1-Fam in 1MEA cells is to the TRP channels, a negative control 10 residue peptide truncated version of Acyl Carrier Protein (ACP) (VQAAIDYING) was chemically synthesized and labeled with FITC (ACP-Fam) ACP-Fam was applied to similar IFA experiments as for Tv1 and truncSorC-Rho with TRPC6 and TRPV6 channels. ACP-Fam was unable to bind to 1MEA cells or colocalize with TRP channels, indicating that colocalization of Tv1-Fam with TRP channels was due to its specific peptide sequence (Fig. 4c).

### Tv1 inhibited COX-2 expression and bioactivity

To determine the intracellular downstream effects of Tv1/TRP channel binding, we examined intracellular molecular mechanisms that are Ca^2+^ dependent and known to be overexpressed in cancer models. COX-2 is overexpressed in many premalignant, malignant, and metastatic cancers, including HCC, and is therapeutically targetable.^36,37^ COX-2 expression is undetectable in most normal tissues, and is highly induced by pro-inflammatory cytokines, mitogens, oncogenes and growth factors. The tumorigenicity of 1MEA cells is associated with human growth factor (HGF) upregulation, which promotes EMT and carcinogenesis via upregulating COX-2 and Akt.^38^ In this study, immunoblot analysis confirmed that COX-2 was overexpressed 2.5-folds in 1MEA cells compared with BNL cells (Figs. 5a and c). Furthermore, when 1MEA cells were treated with Tv1 for 48 h, COX-2 expression decreased by 45% compared with untreated cells (Figs. 5b and d). To determine if Tv1 treatment induced lower COX-2 bioactivity in 1MEA cells, the downstream pathway was further studied by measuring prostaglandin E2 (PGE2) release after Tv1 treatment (Fig. 5e). PGE2 release into the medium of Tv1-treated 1MEA cells decreased by 68.2% (p<0.01) compared with untreated cells. Interestingly, Dox treatment did not lower the PGE2 release by 1MEA cells, suggesting divergent mechanisms of action for Tv1 and Dox.

**Figure 5.**
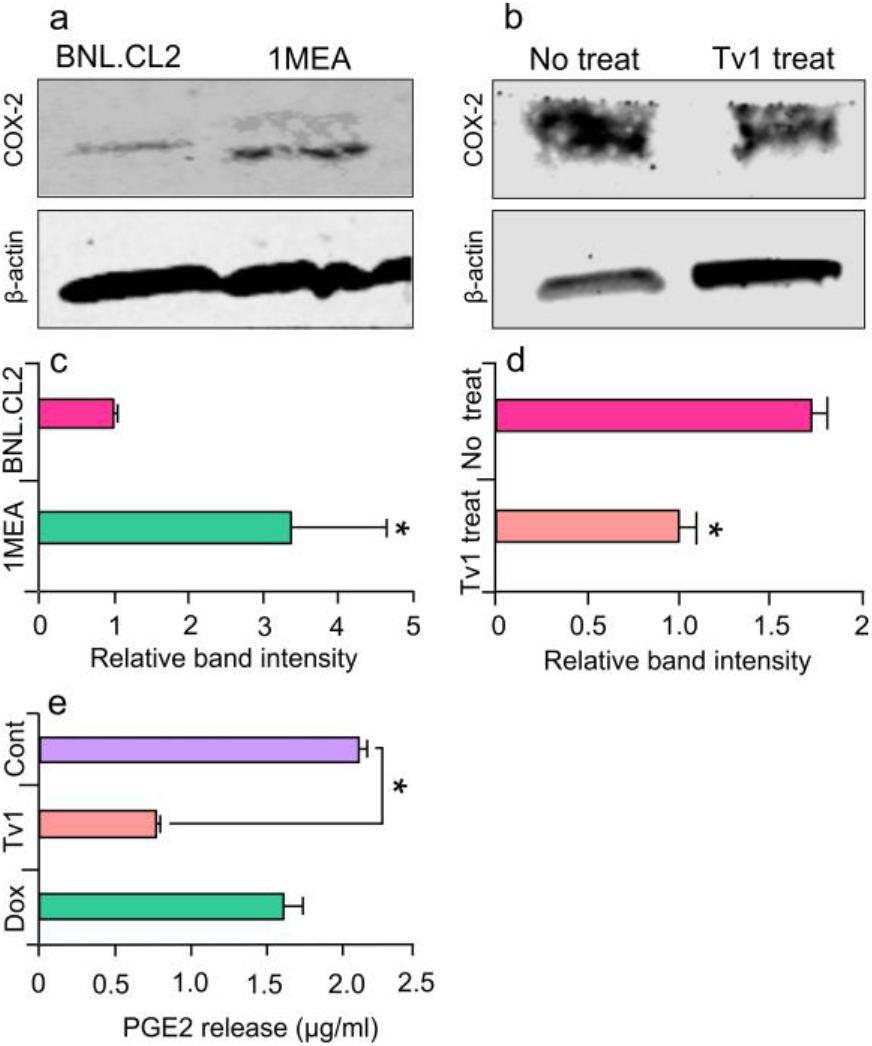
Inhibition of COX-2 expression and PGE2 release by Tv1. **(a)** Immunoblot analysis of COX-2 in normal (BNL.CL2) and cancer (1MEA) cells (n=3). **(b)** COX-2 immunoblot analysis in no treatment and Tv1 treated cells (n=3). (c) Quantitative analysis of bands presenting significant increased levels of COX-2 expression in 1MEA cells when normalized with the ß-actin bands (n=2). **(d)** Significantly lowered expression of COX-2 after Tv1 treatment in 1MEA cells. **(e)** ELISA for prostaglandin E2 (PGE2) release in 1MEA cells after Tv1 treatment showing a significant (p<0.01) decrease in the levels compared with no treatment cells (n=3).

### Molecular modeling of the Tv1/TRPV6 interaction

To further investigate the potential binding of Tv1 to TRPV6 and TRPC6 receptors, we applied molecular modeling techniques using the Cryo-EM structure of human TRPV6 channel (PDB: 6BO8), the solution NMR structure of Tv1 toxin (PDB: 2MIX), and a sequence-built structure for truncSorC, the thirteen residues at the C-terminus of soricidin peptide (Fig. 6).^39^ Our pipeline for modeling applied the Schrodinger software modeling tools ProtPrep and Desmond to prepare the receptor and conduct Tv1 or truncSorC ligand folding and energy minimization, then concluded with molecular dynamic simulations (Fig. S4). Ramachandran plots, RMSD/RMSF data, and RMSD-based trajectory clustering was used to select the most representative ligand structure for Tv1 and SorC models (Figs. S5 & S6). Convincingly, both Tv1 and truncSorC peptides settle into to stable binding modes on the scale of a few hundred nanoseconds. We used the druggability assessment tool SiteMap to find potential general, not-ligand specific, docking sites on the exposed surface of hTRPV6 channel. This mode of Sitemap allows for search of shallow pockets, which is important for peptide binding, as peptide pockets tend to be shallow as a gain of more interactions due to a larger molecule size. Results of SiteMap analysis confirmed that there are several potential favorable Tv1 docking sites exposed on the hTRPV6 extracellular region and in the receptor’s central pore with SiteScores >0.90 and Dscores >0.98 (Fig. 6a). Rigid docking of Tv1 and truncSorC to find potential ligand poses and docking sites was performed using PIPER, and docking results correspond with SiteMap predictions (Fig. 6c & d). To refine docking analysis, clusters of PIPER-derived poses were scored using the Glide/Molecular Mechanics (MM)/Generalized Born Surface Area (GBSA) protocol. Noticeably, hTRPV6’s central pore location gave the lowest energy scores for both Tv1 and truncSorC models (Figs. 6a & 6b). TRPV6’s central pore and adjacent extracellular area were used to build a docking grid and perform flexible docking with GlideSP followed by MM/GBSA analysis. Complexes with the lowest MM/GBSA score, 55.7 for Tv1 and −49.6 for truncSorC were selected for further analysis via molecular dynamics (MD) simulations (Figs. 7a & 7b). Specifically, there were several electrostatic interactions between Tv1 and hTRPV6 at such as, V549, N548, D542, I541, and P544. To validate the resulting trajectory data and binding energy fluctuation over trajectory, 51 frames were uniformly extracted over a 150 ns timeframe and subjected to MM/GBSA calculation (Table S1). The MM/GBSA biding energies were −84.3+9.5 and −61.7+12.2 for Tv1 and truncSorC, respectively, indicating favorable binding conditions. Specifically, there were several Hydrogen bonding interactions between Tv1 and hTRPV6 most notably at residues E519 (A-Chain), D542 (A, B, D – Chains), I541 (B, D-Chains), and N548 (C-chain) of the receptor (Figs. S7-S10). We performed an *in silico* alanine scan mutating to alanine the residues in hTRPV6 that had the strongest interactions and all residues of Tv1 except the cysteine residues and found that the binding affinities changed significantly at the receptor residues E519 (A-Chain), D542 (A, B, D - Chains), and N548 (C-chain), which are also the major sites of receptor-Tv1 interactions (Table 1). Specifically, binding affinities for A:E519, B:D542, and D:N542 experienced a +6.24, +6.27, +12.2 change, respectively. These findings increase the validity of our Tv1 hTRPV6 computational binding model.

**Figure 6.**
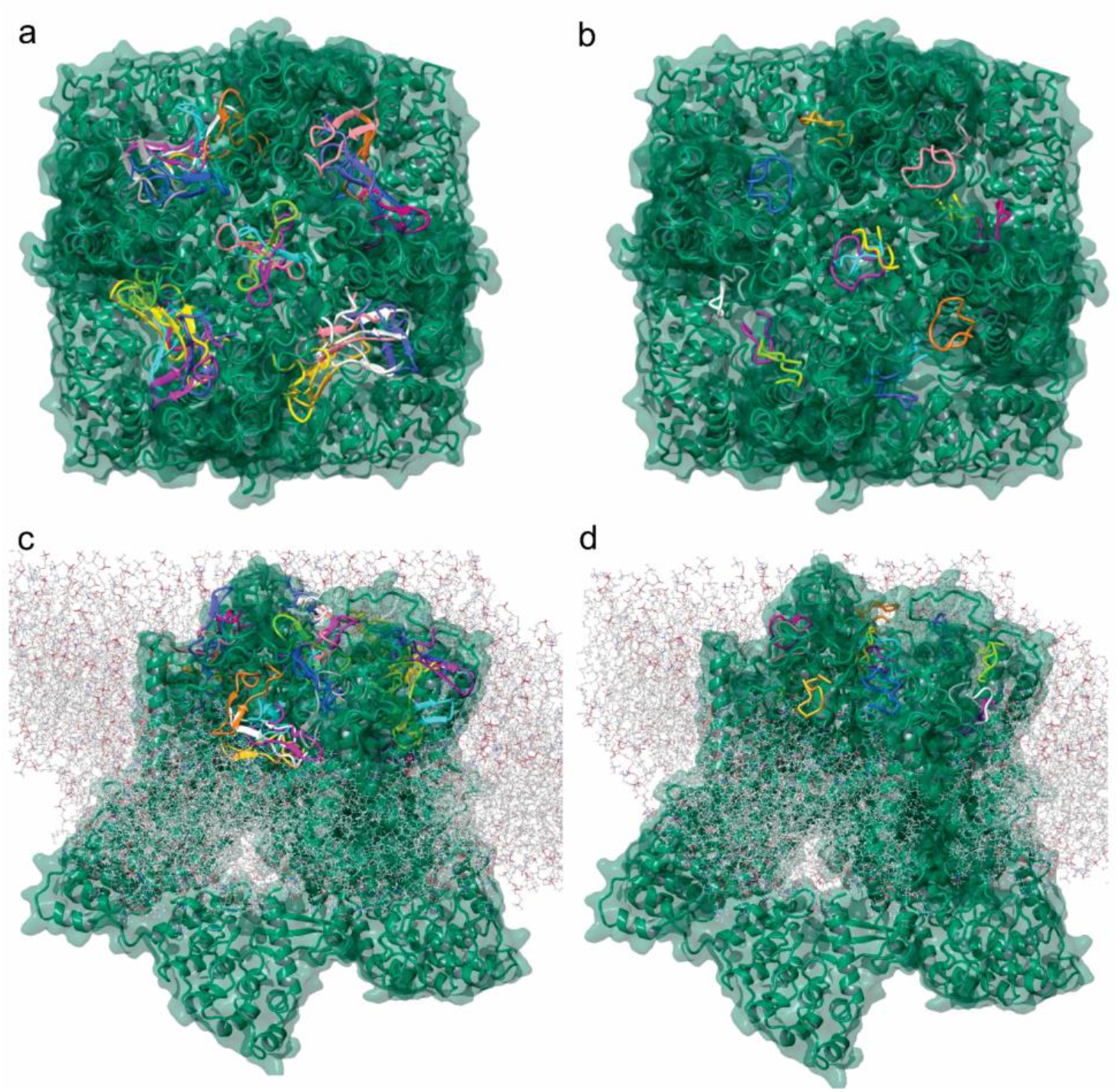
Tv1 and TruncSorC docking to hTRPV6 ion channel. Visualization of potential docking sites of **(a)** Tv1 and **(b)** truncSorC ligands with hTRPV6 calculated by Glide/MM/GBSA docking algorithm. Similar positions as shown for **(c)** Tv1 and **(d)** truncSorC ligand on extracellular surface of hTRPV6 with membrane present using a rigid docking in PIPER algorithm. Visualization with membrane was used to eliminate potential docking sites.

**Figure 7.**
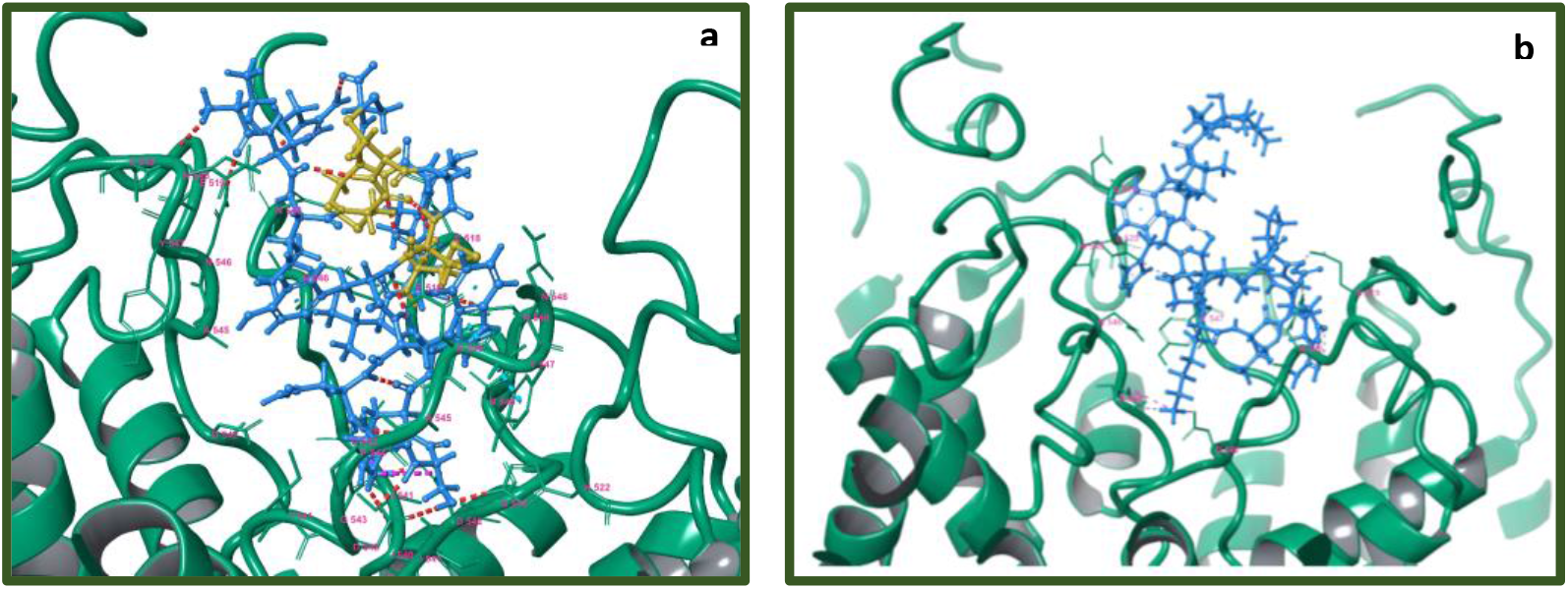
Visualization of lowest energy pose of Tv1 and trunc-SorC with hTRPV6. Receptor represented as teal colored ribbon and its residue in 4Å proximity with (a) Tv1 in blue with cysteine residues in yellow and (b) truncSorC in blue. Poses calculated by Glide/MM/GBSA docking algorithm.

**Table 1.**
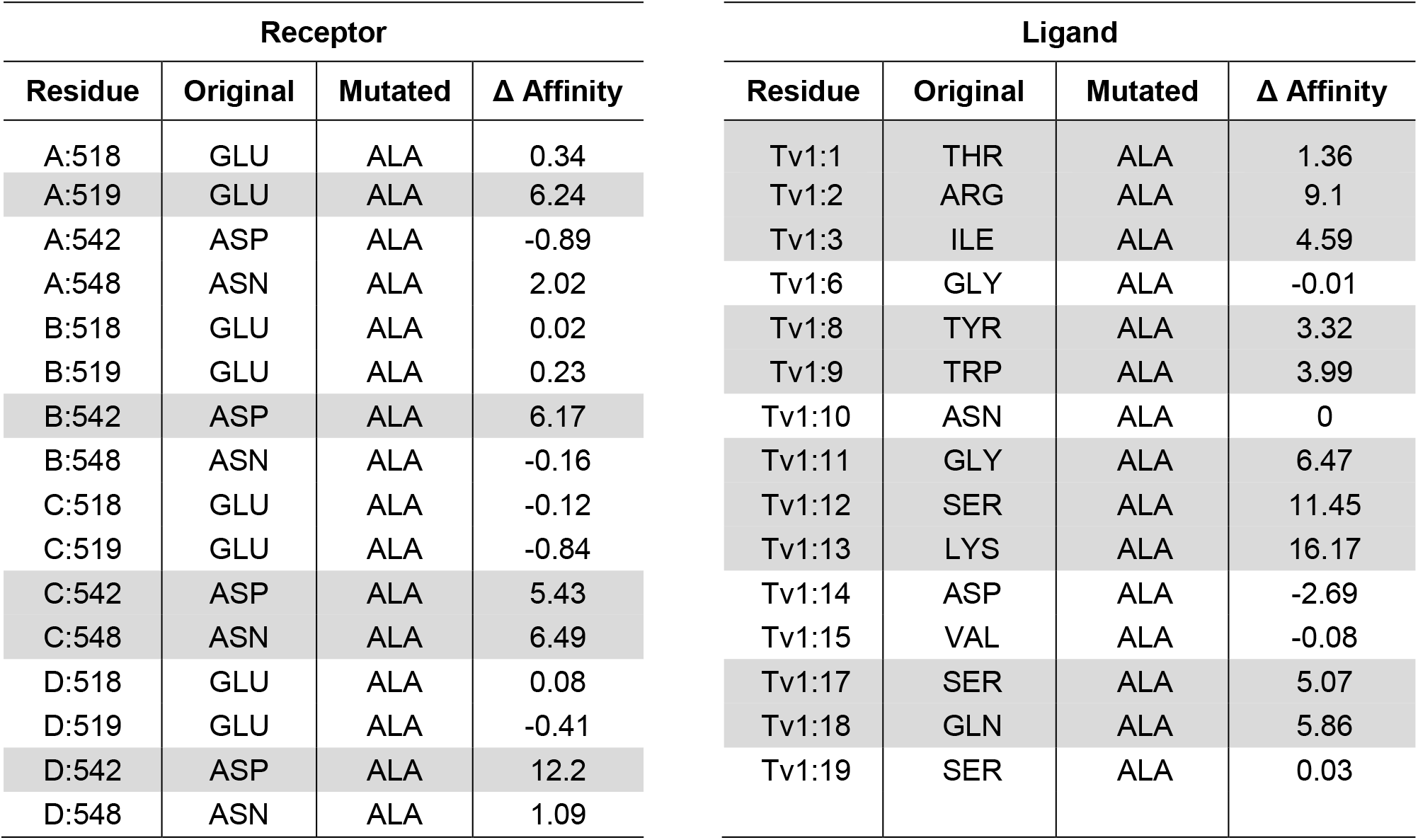
Results of *in-silico* alanine scan of selected hTRPV6 receptor residues and non-cysteine Tv1 ligand

| Receptor |  |  |  |
| --- | --- | --- | --- |
| Residue | Original | Mutated | $\Delta$ Affinity |
| A:518 | GLU | ALA | 0.34 |
| A:519 | GLU | ALA | 6.24 |
| A:542 | ASP | ALA | -0.89 |
| A:548 | ASN | ALA | 2.02 |
| B:518 | GLU | ALA | 0.02 |
| B:519 | GLU | ALA | 0.23 |
| B:542 | ASP | ALA | 6.17 |
| B:548 | ASN | ALA | -0.16 |
| C:518 | GLU | ALA | -0.12 |
| C:519 | GLU | ALA | -0.84 |
| C:542 | ASP | ALA | 5.43 |
| C:548 | ASN | ALA | 6.49 |
| D:518 | GLU | ALA | 0.08 |
| D:519 | GLU | ALA | -0.41 |
| D:542 | ASP | ALA | 12.2 |
| D:548 | ASN | ALA | 1.09 |

| Ligand |  |  |  |
| --- | --- | --- | --- |
| Residue | Original | Mutated | $\Delta$ Affinity |
| Tv1:1 | THR | ALA | 1.36 |
| Tv1:2 | ARG | ALA | 9.1 |
| Tv1:3 | ILE | ALA | 4.59 |
| Tv1:6 | GLY | ALA | -0.01 |
| Tv1:8 | TYR | ALA | 3.32 |
| Tv1:9 | TRP | ALA | 3.99 |
| Tv1:10 | ASN | ALA | 0 |
| Tv1:11 | GLY | ALA | 6.47 |
| Tv1:12 | SER | ALA | 11.45 |
| Tv1:13 | LYS | ALA | 16.17 |
| Tv1:14 | ASP | ALA | -2.69 |
| Tv1:15 | VAL | ALA | -0.08 |
| Tv1:17 | SER | ALA | 5.07 |
| Tv1:18 | GLN | ALA | 5.86 |
| Tv1:19 | SER | ALA | 0.03 |

## Discussion

Bioactive peptides found in the toxic arsenal of venomous organisms are a promising source for anticancer drug therapy as they have the potential to be specific to ion channels that are overexpressed in tumor cells. ^4,40^ In this study, Tv1, a venom peptide from predatory marine snail *T. variegata* showed specific and selective cytotoxicity for murine liver cancer cells by binding to the tumor cell membrane and modulating TRPC6 and TRPV6 ion channels. *In vivo* allograft tumor models demonstrated a potent and significant reduction in tumor size when Tv1 was administered to mouse models, demonstrating that Tv1 is a venom peptide that minimizes the growth of HCC tumor cells.

Tv1’s colocalization with TRPC6 and TRPV6 channels is particularly promising, as TRP channels are known to influence the cell cycle by regulating gene transcription, as well as influencing other cellular processes such as proliferation, apoptosis, and motility.^41^ Many TRPC members are demonstrated to maintain intracellular Ca^2+^ homeostasis and play important roles in cell cycle and Ca^2+^-related factors and pathways. Several studies have shown a close correlation between TRPC6 channel overexpression and the development of cancers, such as prostate, breast, liver, brain, gastric, and oesophageal cancer.^29,42^ TRPC6 contributes to the proliferation of prostate cancer epithelial cells, human epithelial breast cancer cells and human hepatoma cells.^43^ Moreover, blocking TRPC channels leads to significant inhibition of tumor cell proliferation. To date, studies exploring the role of TRPCs in cancer proliferation have mainly focused on TRPC1 and TRPC6, for which spider venom peptide GsMTx4 has been reported to be an allosteric inhibitor.^44,45^

Similar to TRPC6, TRPV6 channel upregulation in prostate cancer cells is known to represent a mechanism for maintaining a higher proliferation rate, increasing cell survival and apoptosis resistance.^46,47^ TRPV6 mRNA and protein expression has been detected in ovarian cancer and other cancer types, such as breast, prostate, thyroid and colon cancer.^35,48^. Due to the involvement of TRPV6 channels in cancer cell proliferation^46,49^ and its overexpression in numerous cancer models, attempts have been made to virtually screen TRPV6’s Ca^2+^ channel inhibitors^50^. The crystal structure for TRPV6 was recently solved and enabled us to model the potential Tv1/TRPV6 interaction (Fig. 6).^51^ TRPV6 is a highly selective Ca^2+^ channel that has been considered part of store-operated calcium entry (SOCE). Furthermore, Ca^2+^ entry through ORAI1-mediated Ca^2+^ channels play a critical role in the migration and metastasis of breast cancer cells. Attenuating SOCE by knocking down ORAI1 protein expression resulted in the reduction of breast cancer cell proliferation.^52,53^ In this study, liver cancer 1MEA cells were also found to overexpress TRPC6 and TRPV6 channels, and immunofluorescence studies showed that Tv1 co-localized with these channels. While TRP channels are widely expressed in different types of cancer cells, here we report the first description of the presence of TRP channels in BNL and 1MEA cells.^54^

Our proposed mechanism for Tv1 anticancer activity via TRPC6 and TRPV6 channels involves the downstream inhibition of the COX-2 pathway (Fig. 1). This mechanism is similar to previous studies investigating colonic myofibroblasts, where NF-κB and NFAT serve as important positive and negative transcriptional regulators of TNF-α-induced COX-2-dependent PGE2 production, downstream of SOCE (Ca^2+^ influx) via TRPC6/V6 channels. ^55^ The inhibitory activity of Tv1 on TRPC6 and/or TRPV6 channels were substantiated by decreased activity of intracellular Ca^2+^-mediated processes. Specifically, the decreased expression of COX-2 and PGE2 release suggested that Tv1 inhibited calcium influx into tumor cells via TRP channels (Fig. 1). Mechanistic studies implicate COX-2 is overexpressed in most solid tumors, including liver, colorectal, pancreatic, breast, lung cancer as well as osteosarcoma.^56,57^ COX-2 promotes angiogenesis, tissue invasion, metastasis, and resistance to apoptosis.^58^ Genetic studies have also supported a cause-effect connection between COX-2 and tumorigenesis. In tumors, COX-2 overexpression leads to increased PGE2 levels, which affect many processes involved in carcinogenesis, such as angiogenesis, inhibition of apoptosis, stimulation of cell growth as well as the increased invasiveness and metastatic potential. Recently it was demonstrated that enhanced COX-2 expression in hepatocytes is sufficient to induce HCC, and its inhibition may provide a potential approach for preventing and treating liver cancer.^36^ COX-2 and PGE2 levels were both significantly decreased following Tv1 treatment in 1MEA cells (Fig. 2b). Experimental studies in animal models of liver cancer have also shown that NSAIDs, including both selective and non-selective COX-2 inhibitors, exert chemopreventive as well as therapeutic effects. However, the key mechanism by which COX-2 inhibitors affect HCC cell growth is not yet fully understood.^56^ While functional electrophysiological assays for Ca+^2^ release in the presence and absence of Tv1 are required to confirm this proposed mechanism of action, our findings suggest Tv1 has downstream intracellular effects on COX-2 and PGE2 activity.

Additionally, Ca+^2^ transport in TRPV6 tetrameric receptor is via the central pore and is regulated by K+ channel-like transmembrane domain.^51^ Analyses of TRPV6 receptor-ligand interactions using virtual docking and molecular dynamics demonstrated that both Tv1 and truncSorC ligand models heavily interact with the extracellular recruitment site formed by E518, E519 and N548 residues and a selectivity filter of the pore region formed by D542 side chains, one from each hTRPV6 subunit.^59^ Our molecular modeling results indicate the energetically favorable position of Tv1 and truncated Soricidin peptide (truncSorC) ligands results in blocking the TRPV6 receptor pore and potentially inhibiting physiological functions pertaining to Ca^2+^ transport. *In-silico* replacement of binding residues by alanine on the hTRPV6 receptor and Tv1 ligand results in loss of affinity between receptor and docked ligand providing an additional understanding about the mode of interactions between Tv1 and hTRPV6 pore (Table 1). While we layered several computational modeling methods together, it is important to highlight the methods, while separate, are all converging on similar results (Fig. 6).

The emergence of ion channels as molecular targets for new cancer therapies has led to immense interest in peptide-based therapies derived from the venom arsenal of predatory organisms.^60,61^ Ion channels and transporters mediate the transport of ions that participate in the regulation of tumor cell survival, death and motility.^62,63^ Increasingly, cancer is being viewed as a channelopathy and as such, new cancer therapeutic targets are aimed at manipulating ion channels and transporters that are differentially expressed in tumor versus non-tumor cells.^64,65^This strategy has significant benefits, as it can address many problems that arise while using conventional cancer therapies, such as tumor cells developing drug resistance, nonspecific toxicity from targeting healthy cells, and the risk for cancer recurrence.^66^ Venom peptides that demonstrate antitumor activity, as shown here for Tv1 from venomous marine snail *T. variegata*, highlight a paradigm shifting approach for advancing efforts to understand and treat cancer.

## Methods

### Tissue culture and reagents

The cell lines BNL1MEA.7R.1 also called 1MEA (mouse liver carcinoma) and BNL.CL.2 (mouse liver epithelium) were a kind gift from Professor Olorenseun Ogunwobi (Hunter College, New York, NY). Cells were maintained in Dulbecco’s modified Eagle’s medium (DMEM, Gibco-BRL Life Technologies, Paisley, UK) supplemented with 10% fetal bovine serum (FBS), 1% penicillin, 4.5g/L glucose, L-glutamine, and sodium pyruvate. HeLa (human cervical cancer), SKnSH (human neuroblastoma), (American Type Culture Collection, Rockville, MD), were regularly cultured in Eagle’s minimal essential medium (Gibco Co., Grand Island, NY) supplemented with 10% fetal bovine serum (FBS), 1% penicillin. RWPE1 (human prostate carcinoma) were cultured in serum-free keratinocyte media containing bovine pituitary extract and epidermal growth factor (GIBCO Invitrogen, Carlsband, CA) at 37°C and 5% CO2. The tetrazolium salt, 3-4, 5 dimethylthiazol-2,5 diphenyl tetrazolium bromide (MTT) and doxorubicin hydrochloride (Dox) were both obtained from Sigma (St. Louis, MO). Sorafenib was a kind gift from Bayer pharmaceutical.

### Synthesis, purification and characterization of Tv1

As teretoxins are present only in minute quantities in the venom, a greater quantity of the linear peptide Tv1 was chemically synthesized and purified by previously described methods and further oxidatively folded into biologically active form^32^ (Figs. S11a & b).

### Fluorescent labeling of oxidized Tv1 peptide

Oxidized Tv1 was non-specifically labeled with 5/6 FAM using NHS ester of fluorescein. Folded Tv1 was dissolved in 0.1M NaHCO3 buffer at a concentration of 1-10μg/ml at pH 8-8.5. For mono labeling of the peptide, 8 times of the NHS-ester was used. Total reaction volume was maintained to 100μl. Reaction was done at 4°C overnight and monitored using UHPLC at 495nm for the fluorescent label and 214nm for peptide bond absorption. Confirmation of the product was done using LC-MS. (Figs. S12a & b)

### Synthesis and Rhodamine labeling of truncated Soricidin (truncSorC) peptide

A known TRPV6 inhibitor truncated soricidin peptide (27 residues long with a sequence of FGK LSSND TEGGL CKEFL HPSKV DLPR) was synthesized by microwave assisted Fmoc solid–phase peptide synthesis on a CEM Liberty synthesizer using standard side chain protection. Following treatment of peptidyl resin with Reagent K [92.5% TFA (Trifluoroacetic acid), 2.5% TIS (Triisopropylsilane), 2.5% EDT (1,2 Ethanedithiol) and 2.5% water, 4 h)] and methyl tertiary butyl ether (MTBE) precipitation, crude checked for purity on Agilent UHPLC machine and eluted using a linear gradient from 0% to 75% buffer B (80% Acetonitrile with 20% water) in 3.5 min. The identity of synthesized peptide was confirmed by molecular mass measurement of purified peptide using 6520 Agilent Q-TOF LC-MS (Figs S13a &b). For colocalization experiment in the presence of Tv1-Fam, the pure peptide was labeled with a different tag Rhodamine NHS ester in this case. Peptide was dissolved in the conjugation buffer at a concentration of (1-10μg/ml) NHS-Rhodamine was dissolved in DMF/DMSO at 10μg/ml. 8 times of the dye was transferred to the peptide and incubated on ice at 4 °C for 2 h. the conjugation reaction was monitored using UHPLC at 555nm for the emission of Rhodamine and 214/280 for peptides (Figs. 14a & b).

### Synthesis of a non-specific fluorescent peptide ACP-Fam

10 residues long a non-specific peptide, a truncated version of acyl carrier protein with a sequence of VQAAIDYING was synthesized as mentioned above and further purified and characterized using Agilent HPLC and LC-MS (Figs. 15a & b). To fluorescently label it, a reaction scheme as shown in figure S16a was applied and the product was characterized by UHPLC and LC-MS (Figs. 16b & c).

### *In vitro* cell viability or MTT Assay

The tetrazolium salt, 3-4, 5 di-methyl thiazol-2, 5 diphenyl tetrazolium bromide (MTT) was obtained from Sigma. MTT was prepared as a stock solution of 5 mg/ml in Phosphate buffered saline (PBS). Dox was prepared as a stock solution of 100mg/ml in deionized water and then further diluted in culture medium to desired concentration. Sorafenib was dissolved in DMSO, 100mM as a stock solution and further diluted in culture medium as desired. All the solutions were filter sterilized before use. Cells were seeded at a density 5000 cells/well, in triplicates, in 96 well plates 24 h before treatment. Following 48 h treatment with Tv1 peptide, Sorafenib and Dox, medium was aspirated from the wells and100ul of 5mg/ml MTT solution was added and incubated for 3 h in incubator at 37°C and 5% CO2. Afterwards, 100μl of 0.04N HCl in isopropanol was added to stop the reaction. Absorbance was measured using a plate reader Gen5 software to analyze for cell viability at 550nm and 620nm and delta values were plotted.

### Flow Cytometry

ApoScreen Annexin V-FITC kit from Southern Biotech was used and protocol was followed according to the manufacturer’s protocol. 1×10^6^ Cells were seeded in 100mm Petridishes 24 h prior to the treatment. To check for early stage apoptosis the cells were treated for 16 h with Tv1. A 4-tube protocol is used to aid in initially setting the correct fluorescence compensation on the flow cytometer. Tubes 1-4 as follows: 1. Unstained cells, 2. Annexin V-FITC only, 3. PI (propidium iodide) only, 4. Annexin V-FITC+PI. Cells were washed twice in cold PBS and resuspended in cold 1X binding buffer to a concentration of 1 × 10^6^ to 1 × 10^7^ cells/mL. 100 μL of cells (1 × 10^5^ to 1 × 10^6^) were added to each tube. Add 10 μL of Annexin V-FITC was added to tube 2 and tube 4, gently vortexed and incubated for 15 minutes on ice, protected from light. 380 μL of cold 1X binding buffer was added to each tube without washing. And now, 10 μL of PI was added to tube 3 and tube 4 without washing. FACS Calibur Flow Cytometer (BD Biosciences) was used to analyze the samples.

### Migration Assay

1 × 10^4^ cells were seeded into 6-well plates and at 70% confluency, the cell monolayer was wounded with a 200μl-pipete tip, washed with PBS and medium was replaced with peptide/drug containing medium. Images were taken at different time points at time 0 and then further at 4, 6, 12 and 24 h later. Images from 3 experiments were analyzed for percentage of cell-covered area using the Wimasis Image Analysis software (Wimasis GmbH, Munich, Germany).

### Antitumor activity in allograft transplantation models

All animal work was carried out in accordance with the general public health service guidelines (PHS policy at IV.A.3.B) and approved by Institutional Animal Care and Use Committee (IACUC) at Hunter College with a protocol #2015-0038.

Cancer immunotherapies are designed to work in conjunction with a patient’s immune system to increase native anti-tumor responses. In this field of study, conventional xenograft models lack relevance due to the animals’ immunocompromised status. A syngeneic mouse model, however, provides an effective approach for studying how cancer therapies perform in the presence of a functional immune system.^67^ To evaluate the in vivo antitumor activity of Tv1, a syngeneic hepatocellular carcinoma model in mice was used. It was established by subcutaneous injection of 5×10^6^ BNL 1ME A.7R.1 cells into the left flank of female Balb/c mice. Once the BNL 1ME A.7R.1 allografts reached a size of ~250 mm^3^, fifteen mice were randomly assigned to one of the three groups. In two of the three groups (group 1 and 2), compound Tv1 was administered intraperitoneally at a dose of 0.08 and 0.8 mg/kg body weight (mpk) respectively while the third group was used as a control. Tv1, as well as the vehicle control, were administered daily for eight days and a four days break was given before starting the treatment again for seven days. Tumor volumes, and body weights were recorded every alternate day, until the termination of treatment.

### In cell western assay

To screen the presence of different subtypes of TRP (transient receptor potential) channels on the cells, in-cell western technique was used. BNL.CL2 and BNL.1MEA.7R.1 cells were seeded in 96 well plates with a density of 1×10^4^ per well and cultured overnight. On Day 2, the cells were fixed in 4% PFA (paraformaldehyde) by adding 20 μl of 12% PFA directly to the wells for 1 h at room temperature. The wells were washed three times with PBS (50 μl/well), permeabilized with PBS/0.1% Triton X-100 (50 μl/well, three times, 2 mins each), and blocked in LI-COR buffer (50 μl/well) for 2 h at room temperature (or alternatively overnight at 4°C). Assigned wells were then incubated with mouse HERG, and different TRP channel subtype antibodies (1: 200 for optimal signal-to-noise ratio) in LI-COR blocking buffer for overnight in cold room (20 μl/well) and next day washed with PBS/0.1% Tween-20 (50 μl/well, three times). Infrared anti-mouse IRDye800CW secondary antibody (1: 1,000) and cell tag (1: 500) in Licor blocking buffer with 0.5% Tween-20 were then added (50 μl/well). The plates were incubated for 1 h at room temperature, and the wells were washed with PBS/0.1% Tween-20 (three times, 5 min each). The plates were covered with black seals and imaged on an Odyssey infrared scanner using microplate2 settings with sensitivity of 5 in both the 700 and 800 nm wavelength channels. Data were acquired by using Odyssey software, exported and analyzed in Microsoft Excel. Cell tag values were background subtracted from wells treated only with secondary antibody, and then normalized to cell numbers by dividing by the total 800 nm signal.

### Immunofluorescence Assay

After 24 h of cells grown on coverslip in 4 well dishes, were incubated with either Tv1-Fam or as indicated in results section for different experiments. After the treatment, cells were fixed in cold methanol at −20 °C for 20 min, washed with PBS, and permeabilized in 0.1% Triton X-100 for 5 min, three times and washed with PBS. After incubation in blocking solution (5% BSA, 1% Goat serum in PBST) for 1 h, cells were incubated with primary antibodies (anti-rabbit polyclonal TRPC6 or TRPV6) in blocking solution for overnight at 4 °C, washed and incubated with secondary antibodies for 1.5 h the next day. Coverslips were mounted on slides using Antifade mounting medium containing DAPI for nuclear staining.

### Confocal Imaging and Colocalization Analyses

For colocalization analyses, the cells were first incubated with the active peptide Tv1 Fam or the known channel blocker (truncSorC-Rho) and further immunostained for the channels. Imaging was done using Nikon A1 Confocal microscope equipped with the Nikon elements acquisition software. Image processing (cropping, contrast adjustment, and background subtraction) and analysis were performed NIS elements. Colocalization coefficients were computed using the same.

### Manders overlap coefficient

Manders Overlap Coefficient (MOC), was implemented in image analysis with the software packages NIS elements.^68^ The Mander’s overlap coefficient (MOC) is used to quantify the degree of colocalization between fluorophores. The MOC was introduced to overcome perceived problems with the Pearsons correlation coefficient (PCC). The two coefficients are mathematically similar, differing in the use of either the absolute intensities (MOC) or of the deviation from the mean (PCC). MOC values are never negative and it’s value is independent of signal levels of both probes and is sensitive to occurrence in the same pixel. MOC is 0 when both probes are completely exclusive. The relevance of this coefficient is that it is possible to obtain the fraction of one probe overlapping with the other and vice versa.

### Whole-cell protein extraction and immunoblotting

Cells were grown in 10% FBS-containing medium as a monolayer in 75cm^2^ flasks until confluent, treated or not treated based on the experiment, cells were washed twice in 10ml of cold PBS and then incubated in cold 1ml RIPA lysis buffer (Amresco) per 5×10^6^ cells on ice for 10 min. Incubated on ice for 15 min, occasionally swirling the flask to keep the surface evenly covered. Cell scraper was used to harvest the cells from the flask. Cell lysate was passed through a pipette several times to form a homogenous lysate and centrifuged at 14,000 × g for 15 minutes at 4°C to separate the total protein (supernatant) from the cellular debris (pellet). Supernatants were transferred to a new microcentrifuge tube on ice. Proteins were either used immediately or stored frozen at −20°C until needed. Protein concentration was quantified using the Bio-Rad Bradford protein assay with bovine serum albumin as standard. 20 μg of protein were loaded onto gradient gels (4-15%) purchased from BioRad and transferred onto PVDF (polyvinylidene difluoride) membrane. After blocking the membranes with 5% BSA in Tween 20, blots were incubated overnight in a cold room in 1:200 COX-2 (mouse) polyclonal antibody (aa 570-598) from Cayman chemical or 1:1000 monoclonal anti-β-actin antibody produced in mouse (Sigma Aldrich). To detect the specific protein bands, IRDye^®^ 800CW goat anti mouse secondary antibody was used in blocking buffer at room temperature on an orbital shaker for 2 h, after washing the membranes 3 times for 10 min with wash buffer (PBS-Tween), blots were scanned with Licor Odyssey^®^ CLx Imaging System. Bands were quantified using the same program and average of 3 blots was plotted in the relative band intensity graph.

### Prostaglandin E2 (PGE2) release

1×10^3^ 1MEA cells per well were seeded into 96 well plates, after 24 h cells were treated with Tv1 for 48 h and untreated cells were used as control. Analysis of PGE2 secreted into serum-free medium was performed as previously described.^69^

### Model preparation and docking

All models were prepared using the Protein Preparation Wizard (Maestro v 11.4). Bond order and formal charges were assigned, and hydrogen atoms were added. To further refine the structure, an OPLS3 force field parameter was used to alleviate steric clashes and the minimization was terminated when heavy atoms RMSD reached a maximum cutoff value of 0.30 Å. Protonation states were assigned to residues according to the pKa based on pH = 7.0 using the Epik v4.2 module. Protein–protein interaction (PPI) specific SiteMap mode was used to detect shallow binding sites by decreasing the amount of enclosure and the threshold for van der Waals interactions (to 0.4 and 0.55).^70^ Sites were kept if they comprised at least 15 site points, a restrictive hydrophobicity definition, a standard grid (1.0 A°) were used. The following properties of the binding sites were calculated by the SiteMap program: size, volume, degree of enclosure/exposure, degree of contact, hydrophobic/-philic character, hydrophobic/-philic balance and hydrogen-bonding possibilities (acceptors/donors). SiteScore and Dscore were derived as

SiteScore = 0.0733 sqrt(n) + 0.6688 e − 0.20 p

Dscore = 0.094 sqrt(n) + 0.60 e − 0.324 p

Where n is the number of site points (capped at 100), e is the enclosure score, and p is the hydrophilic score, and is capped at 1.0 for SiteScore to limit the impact of hydrophilicity in charged and highly polar sites. Dscore uses the same properties as SiteScore but different coefficients and the hydrophilic score for Dscore is not capped.

PIPER was set to use 70000 ligand rotations to probe and the top 1000 results of rigid docking of the ligand were clustered do get a docking pose. Constrains were used to discourage docking in the transmembrane and intracellular regions based on TRPV6 OPM data.

Glide algorithm optimized for polypeptides was applied to build receptor grids set to centroids of PIPER derived docking regions and allowing to fully accommodate peptide ligands (Tv1, truncated Sor C) GlideScore (Gscore) and MM-GBSA score were derived to select best docking model.^71,72^

### Molecular dynamics (MD) simulation

For molecular dynamics simulations, each structure was placed in a cubic cell, using DESMOND v 5.2 System Builder workflow, with size adjusted to maintain a minimum distance of 10 Å to the cell boundary, SPC water was added with an appropriate number of ions to establish 0.15M NaCl concentration. All molecular dynamics simulations were completed using DESMOND v5.2 package. The equations of motion were integrated using the multistep RESPA integrator with an inner time step of 2.0 fs for bonded interactions and non-bonded interactions within the short range cutoff. An outer time step of 6.0 fs was used for non-bonded interactions beyond the cutoff. A Nose–Hoover thermostat with relaxation time of 1.0 ps was utilized to maintain the constant simulation temperature and the Martyna–Tobias–Klein method with relaxation time of 2.0 ps was used to control the pressure. Smooth particle-mesh Ewald method with tolerance of 1e-09 was used to calculate long-range electrostatic interactions. Short range electrostatic interactions were calculated with cutoff radius at 9.0Å. The system was equilibrated with the default protocol provided in DESMOND v5.2, which consists of steepest descent (SD) minimization with a maximum of 2000 steps and a gradient threshold of 50 kcal/mol/Å, followed by 12 ps of Berendsen NVT (constant number of particles, volume, and temperature) simulation at 10 K, followed by 24 ps of Berendsen NVT (constant number of particles, pressure, and temperature) equilibration at 300 K. Default equilibration was followed by a 300ns minimization run for Tv1 and truncated Sor C ligand models and 150ns production run using Glide-derived receptor ligand complexes embedded into POPC membrane. All runs used NTP ensemble at 300 K. Energy was saved in 1.2 ps intervals and trajectory was saved in 30 ps intervals. In order to verify our docking model we performed in silico, alanine scan using Bioluminate package on a clustered representative MD frame.^73^ All non-cysteine residues of Tv1 ligand and four binding residues on a receptor side (E518, E519, N548 and D542) were substituted by alanine and scored using MMGBSA (Table 1).

## Acknowledgments

MH acknowledges funding from the Camille and Henry Dreyfus Teacher-Scholar Award and NSF awards CHE-1247550 and NIH-NIMHD grant MD007599. Work in Dr. Ogunwobi’s laboratory is supported by the NIMHD/NIH grant to Hunter College: 8 G 12 MD007599.

## Author contributions

*Conceptualization*: PA, MH; *Investigation*: PA, PF, JH, ML, MH, KH; *Data Analysis*: PA, MH, OOO; *Writing:* PA, MH and OOO; *Visualization:* PA, MH, PF, JH, ML, MH, CS, KH, OOO; *Supervision:* MH and OOO; *Funding Acquisition:* MH and OOO.

## Additional information

Supplementary Information accompanies this paper at……

## Competing interests

The authors declare no competing interest.

